# Collaborative Cross Founder Expression Analysis (CCFEA)

**DOI:** 10.1101/2021.03.04.422591

**Authors:** Richard R. Green, Reneé C. Ireton, Martin Ferris, Kathleen Muenzen, David R. Crosslin, Michael Gale

## Abstract

To understand the role of host genetic variation in infection by SARS-CoV and influenza A virus we developed the Collaborative Cross Founder Expression Analysis (CCFEA), a shiny visualization tool using public RNAseq data from the collaborative cross (CC) founder strains (A/J, C57BL/6J, 129s1/SvImJ, NOD/ShILtJ, NZO/HILtJ, CAST/EiJ, PWK/PhJ, and WSB/EiJ) that underwent infection by either virus, linked with genetic analyses to define loci linked to infection, immunity, and disease phenotypes. Individual gene expression data is displayed across founders, viral infections and days post infection.

**Resource Table:** 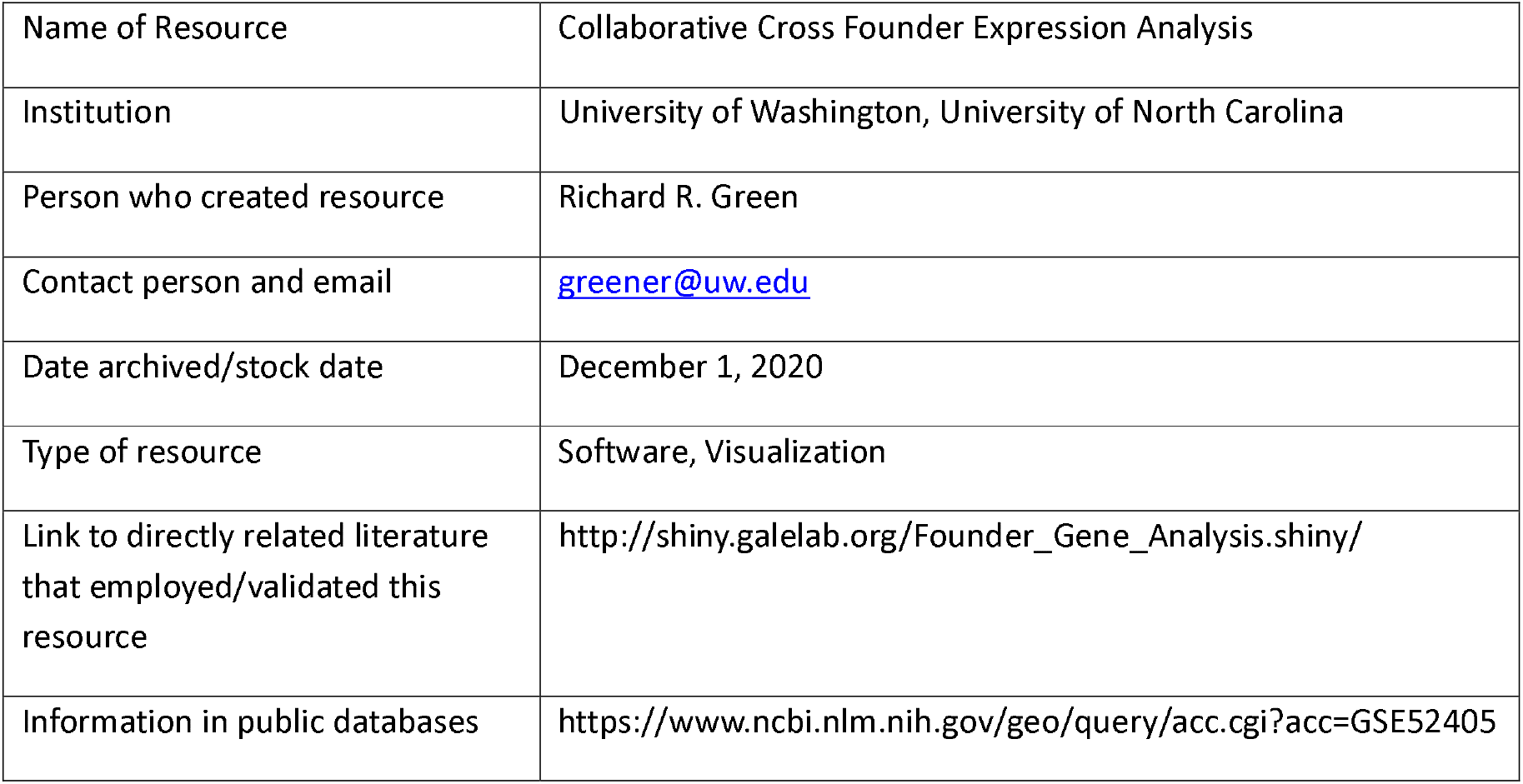

## Resource Details

The Collaborative cross (CC) is a mouse genetics project (http://csbio.unc.edu/CCstatus) using a genetic reference population (GRP) derived from eight (3 wild-derived and 5 inbred) mouse founder strains (A/J, C57BL/6J, 129s1/SvImJ, NOD/ShILtJ, NZO/HILtJ, CAST/EiJ, PWK/PhJ, and WSB/EiJ). Through this complex breeding, the collaborative cross produces hundreds of independently bred, octo-parental recombinant inbred mouse lines (RIX). These RIXs possess complex genetic back grounds that produce diverse phenotypic responses to viral infection (Ferris et al. 2013, Gralinski et al. 2015, Green et al. 2017, Green et. al 2016, Graham et al. 2016, Rasmussen et al. 2014). These studies revealed genome coordinates that were influenced by infection phenotype, similar to genome wide association studies (GWAS). These genetic regions (loci) frequently contain multiple genes but lack an unbiased approach to determine their individual gene contributions. CCFEA calculates the normalized gene expression contributions across founder strains.

**Figure 1.**
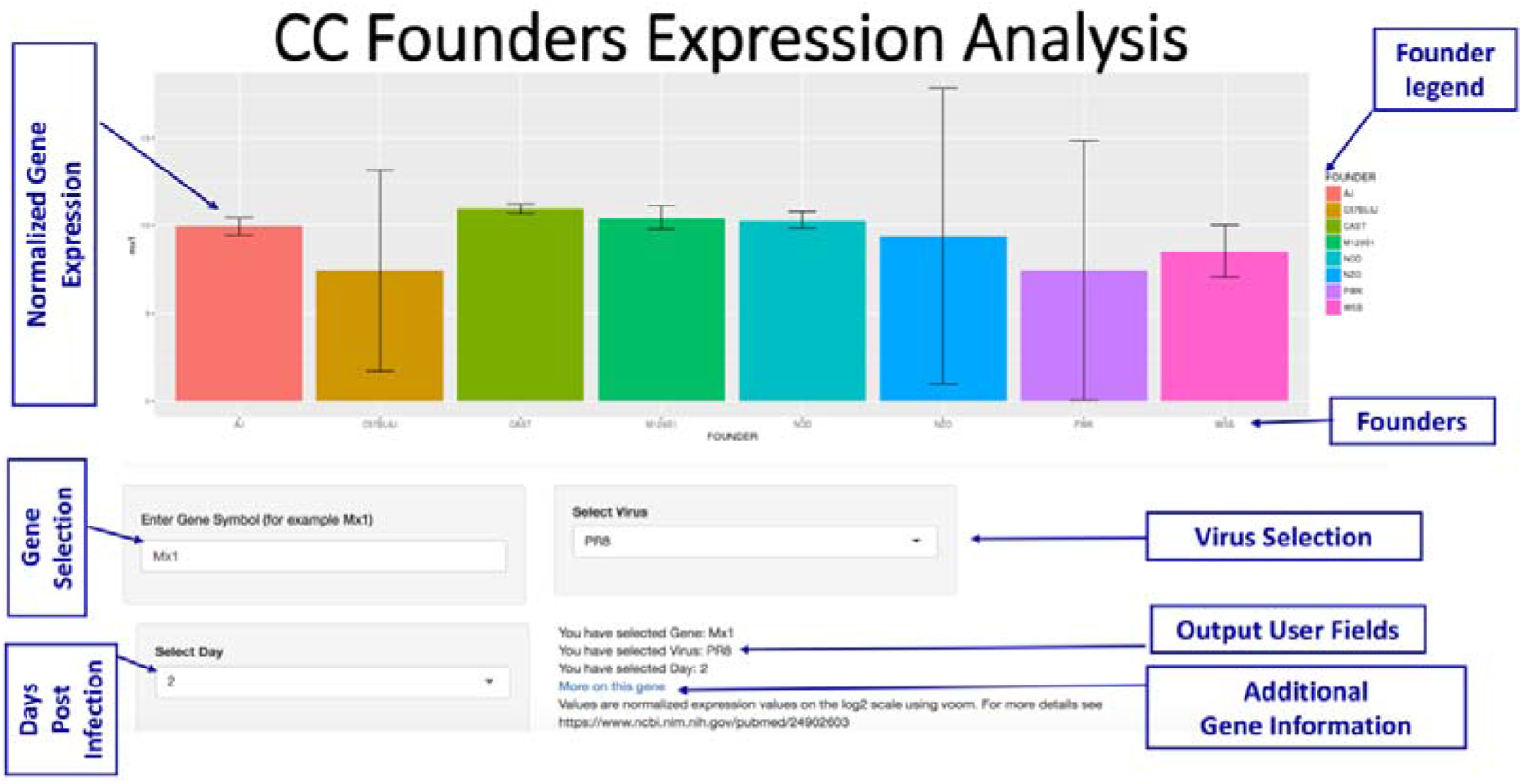
An overview of CCFEA features. We created CCFEA to answer distinct questions about specific gene expression and disease in the CC founders. The application was built in the R statistical programming language and uses the shiny package to query gene intensity across all eight collaborative cross founder strains. Error bars show the natural variation displayed between replicates. Users can select gene signatures across a time course under different experimental conditions which include Influenza A virus infection, severe acute respiratory syndrome coronavirus (SARS-Cov) infection. The Hao et. al study assessed infection responses pathways in tissues but did not address distinct gene expression differences during infection across CC founders.

**Figure 2.**
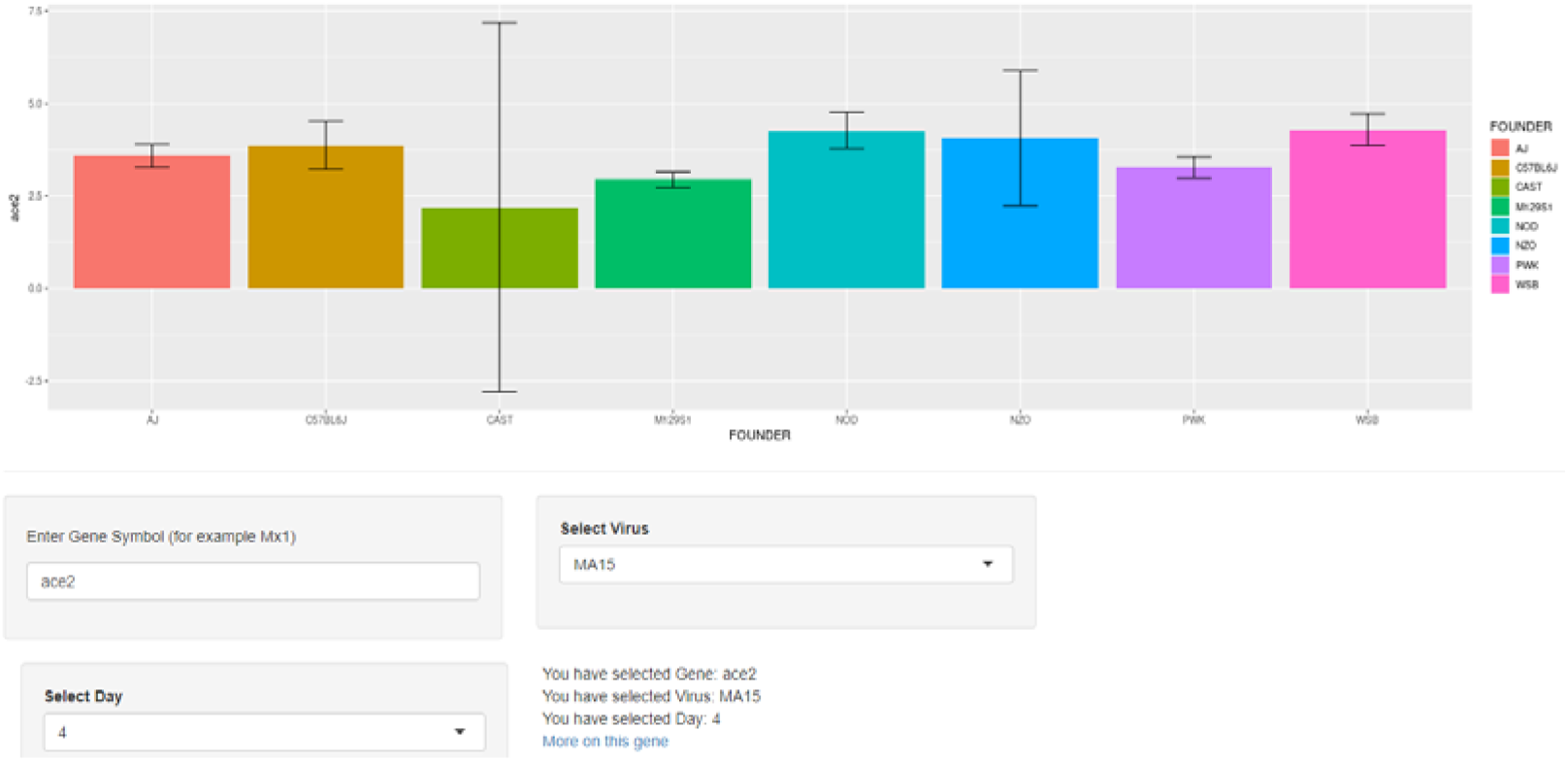
ACE2 mapping in Lung Tissue at Day 4 post infection of SARS-CoV. CCFEA provides valuable insights into the role genetics plays during infection by the betacoronavirus, SARS-CoV, a virus similar to the pandemic virus SARS-COV-2. Both viruses use the host receptor protein ACE2 for cell entry. ACE2 undergoes different baseline expression changes during infection (Figure 2) and recently has been tied to COV19 disease pathogenesis (Bourgonje et al. 2020). Figure 2 shows differences in mapped RNA sequencing reads for the gene encoding ACE2. Some of the mapping differences are influenced by SNPs across the different genetic backgrounds. Differences in the Cast background are possibly influenced by a potential cis regulatory variant. Overall expression changes could be influenced by differences in cellular composition within the infected lungs over time.

## Materials and Methods

RNA was prepared and sequenced (Total RNAseq) with methods described in the Xiong et al. study. This data was also used in a separate study by Josset et al. 2014. The Xiong study addressed the identification of novel transcripts and differential isoforms. The Josset study looked at noncoding expression. CCFEA visualizes the normalized gene counts from the Xiong et al. 2014 study and does not cover the non-coding expression. For CCFEA, raw RNAseq count data was normalized and converted to log counts per million (log cpm) using the voom function in the limma Bioconductor package (Law et al. 2014). Our shiny application features a simple UI that customizes visual output. Shiny is an open source package that creates interactive web applications. CCFEA was built with R version 3.2.5. More information about Shiny can be found here: https://shiny.rstudio.com/.

ARCHS4 is an online resource built off public mouse and human RNAseq datasets (Lachmann et al. 2018). The results include associate genes and pathways that share co-expression patterns. It also provides functional results including predicted biological functions, transcription factors, protein-protein interactions, and expression signatures within tissue types. ARCHS4 was built and maintained by the Ma’ayan lab in the Mount Sinai center for bioinformatics. Additional gene information is provided by ARCHS4 and can be found here: (https://amp.pharm.mssm.edu/archs4/).

## Conclusion

RNA sequencing has become the standard for transcriptional studies due to its speed, cost, and sensitivity. Hosting expression data within shiny in CCFEA allows users to rapidly mine large expression data sets from the R statistical programming but with a simple HTML user interface and does not require any coding experience or training. Shiny is a robust platform frequently used in the biomedical sciences. (Brink et al. 2018, Zhang et al. 2018).

## Acknowledgements

Supported by National Institutes of Health grants U19AI100625, R01AI104002, and U19AI083019. The funders had no role in the design, data acquisition, analysis, or preparation of the manuscript. Thanks to Lucy (Logan) Green for proof reading. Thanks to Gail Jarvik for technical support and suggestions. Thanks to Darla Miller and Ginger Shaw for their support with mouse resources.

